# GBStools: A Unified Approach for Reduced Representation Sequencing and Genotyping

**DOI:** 10.1101/030494

**Authors:** Thomas F Cooke, Muh-Ching Yee, Marina Muzzio, Alexandra Sockell, Ryan Bell, Omar E Cornejo, Joanna L. Kelley, Graciela Bailliet, Claudio M. Bravi, Carlos D Bustamante, Eimear E Kenny

## Abstract

Reduced representation sequencing methods such as genotyping-by-sequencing (GBS) enable low-cost measurement of genetic variation without the need for a reference genome assembly. These methods are widely used in genetic mapping and population genetics studies, especially with non-model organisms. Variant calling error rates, however, are higher in GBS than in standard sequencing, in particular due to restriction site polymorphisms, and few computational tools exist that specifically model and correct these errors. We developed a statistical method to remove errors caused by restriction site polymorphisms, implemented in the software package GBStools. We evaluated it in several simulated data sets, varying in number of samples, mean coverage and population mutation rate, and in two empirical human data sets (N = 8 and N = 63 samples). In our simulations, GBStools improved genotype accuracy more than commonly used filters such as Hardy-Weinberg equilibrium p-values. GBStools is most effective at removing genotype errors in data sets over 100 samples when coverage is 40X or higher, and the improvement is most pronounced in species with high genomic diversity. We also demonstrate the utility of GBS and GBStools for human population genetic inference in Argentine populations and reveal widely varying individual ancestry proportions and an excess of singletons, consistent with recent population growth.

## Author Summary

Eukaryotic genomes range from millions to billions of base pairs in size, but for many genetic experiments it is sufficient to gather information from just a fraction of these sites. In practice, selecting a consistent set of sites can be achieved by cutting genomic DNA with enzymes that recognize DNA sequence motifs, and then sequencing the ends of the resulting fragments. The advantages of this well-known approach are its low cost relative to whole-genome sequencing (WGS), and that it does not require a sequenced genome. These methods, for example genotyping-by-sequencing (GBS), are popular for mapping genes and studying population genetics, particularly in non-model organisms. Here we demonstrate, however, that computational tools designed for WGS are insufficient for handling certain error types that arise in GBS and other similar methods. We present a modified protocol for GBS and a statistical method for detecting these errors, implemented in the software package GBStools. We tested our methods on human DNA samples from Argentine populations. Our results reveal widely varying degrees of European and Native American ancestry, and that rare genetic variants are more numerous than would be expected in a population with constant size.

## Introduction

High-throughput reduced-representation sequencing methods[1] are inexpensive, suffer little from ascertainment bias, and generate genetic markers that are approximately randomly distributed throughout the genome. These methods have been successfully used in trait mapping[2,3], linkage map construction[1,4], selection scans[5,6], and estimating genetic diversity[7]. One such method is genotyping-by-sequencing[8] (GBS). In GBS, the sequencing target is reduced to < 5% of the genome by ligating sequencing adapters only to restriction enzyme cut sites (Fig. 1A). GBS reads can also be assembled into short contigs, which enables single nucleotide variant (SNV) calling without the aid of a genome sequence[9]. Hence, GBS is a popular approach in non-model systems, which typically lack resources such as genome assemblies and microarrays.

**Fig. 1.**
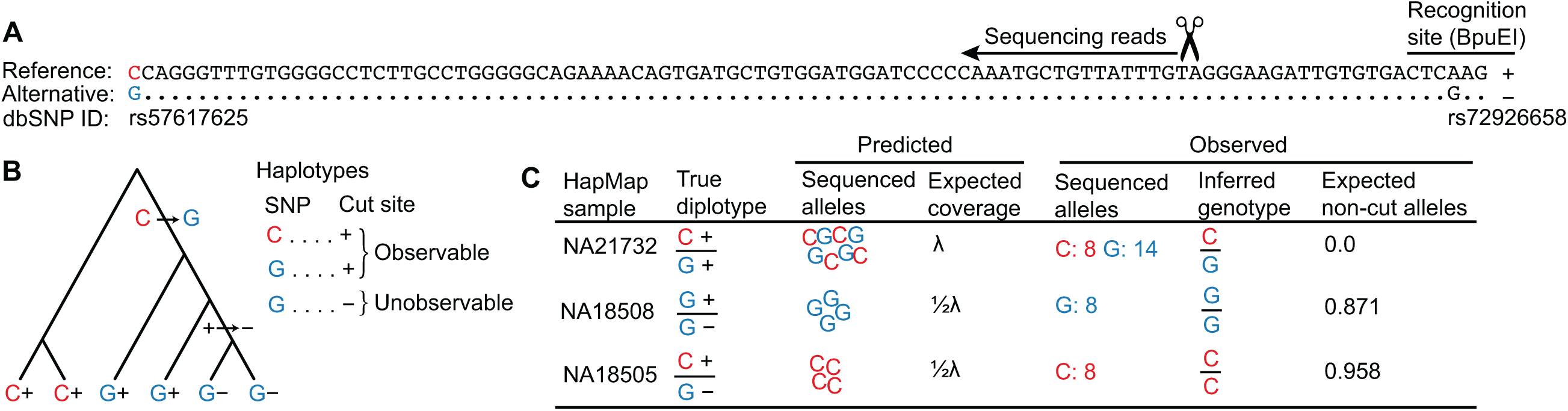
Incorrect inference of genotypes due to restriction site polymorphism. **A.** GBS reads spanning the SNP rs57617625 originated from a polymorphic BpuEl site 94 bp upstream. The non-cut BpuEI allele caused by SNP rs72926658 is labelled as ‘−’ and the cut allele ‘+’. **B.** The ‘−’ allele arose on the haplotype with the derived G allele, causing some G alleles to be unobservable by GBS. **C.** The samples shown carried the three possible heterozygous diplotypes. The sequencing results were consistent with the predictions. Sample NA18505 was incorrectly called homozygous, but the expected non-cut allele count calculated by GBStools (0.958) closely matched the true count (1), identifying it as a probable mis-call.

Unlike whole genome sequencing (WGS), GBS is prone to variant calling errors due to restriction site polymorphisms[7,10–13] (‘allelic dropout’, Fig. 1B). Allelic dropout in GBS can confound applications that rely on accurate calling of rare variation, such site frequency spectrum estimation in population genetics. Here, we present a modified GBS protocol, similar to ddRAD-seq[14], and quantify its error rate. In addition, we present a systematic statistical approach to detect allelic dropout in GBS sequence data, implemented in the open-source software package GBStools.

This approach is based on the fact that allelic dropout reduces a sample’s read coverage at a particular site in proportion to the number of non-cut restriction site alleles it carries there (Fig. 1C). Therefore GBStools models coverage of each sample at a particular site as an overdispersed Poisson random variable drawn from either a distribution with mean λ (zero non-cut alleles carried), a distribution with mean ½λ (one non-cut allele), or with mean zero (two non-cut alleles). GBStools calculates the maximum-likelihood estimate of the parameter λ by expectation-maximization (EM), with the true number of non-cut alleles per sample serving as latent (unobserved) variables (S1 Appendix). The expected values of these latent variables can be used to estimate which samples carry a non-cut allele (see “Expected non-cut alleles” in Fig. 1C). Simultaneously, GBStools estimates the site frequency of the observable reference and alternative SNP alleles, ϕ_1_ and ϕ_2_ (for example see Fig. 1B), and the non-cut allele, ϕ_3_, where ϕ_1_ + ϕ_2_ + ϕ_3_ = 1. Finally, it performs a likelihood ratio test comparing the null hypothesis ϕ_3_ = 0 to the alternative hypothesis ϕ_3_ > 0. In its current implementation GBStools cannot infer the true genotypes obscured by allelic dropout, but it can be used to remove errors by filtering out sites where a high likelihood ratio indicates the presence of restriction site polymorphism.

Lastly, we describe the application of these methods to an extant mixed ancestry population from Argentina to test the performance of GBS in ancestry estimation and demographic inference.

## Results and Discussion

We estimated the magnitude of GBS errors caused by restriction site polymorphisms from both simulated and real data. We chose human as a model system for GBS methods development due to the availability of a high-quality reference genome assembly, high-coverage whole-genome sequencing data,[15,16] and dense SNP array data.

First, we prepared modified GBS libraries from eight HapMap samples from a diverse range of populations and sequenced them on a single HiSeq lane (S1 Table, methods). We used the methylation-insensitive enzymes BpuEI, BsaXI, and CspCI, which cut away from their recognition site. Although a well-balanced mix of different sequencing adapters is commonly used to ensure that restriction enzyme recognition sequences are not over-represented at the start of the sequencing reads [3,4,8,14], our method tolerates low-diversity mixes of adapters, which is convenient when working with smaller sample sets. We quantified each sample by bioanalyzer after PCR, but before pooling, with the goal of reducing variance in the number of reads per sample in the final library. We found, however, that errors at this stage, particularly those caused by incorrect quantification of the bioanalyzer internal standard, can in fact lead to the opposite effect (S1 Fig.). More careful quantification by bioanalyzer, or quantification by fluorimetry, should correct this problem and lead to the desired effect. The HapMap samples had 16.7X mean coverage in a 128 Mb target region (S2 Fig. A, S3 Fig. A). We used GATK[17] to call SNPs in the target regions, and found 483,381 segregating sites that passed variant quality score recalibration. After applying hard filters (coverage ≥ 8X in 8/8 samples, mapping quality ≥ 30, SNP quality ≥ 30), these GBS genotype calls were 98.0% concordant with heterozygous calls from whole-genome sequencing data gathered from the same set of samples (Fig 2A, S2 Table A). We found the error rate dropped as sequencing coverage increased up to 30-40X, after which further increases in coverage had little effect (Fig. 2B). Furthermore, the error rate for singletons was roughly two-fold higher than for non-singletons (Fig. 2C). A filter for known restriction site polymorphisms in the 1000 Genomes Project[18] data set also had a strong effect on concordance (Fig. 2A-C). These three factors appeared to be the major determinants of genotype calling accuracy.

**Fig. 2.**
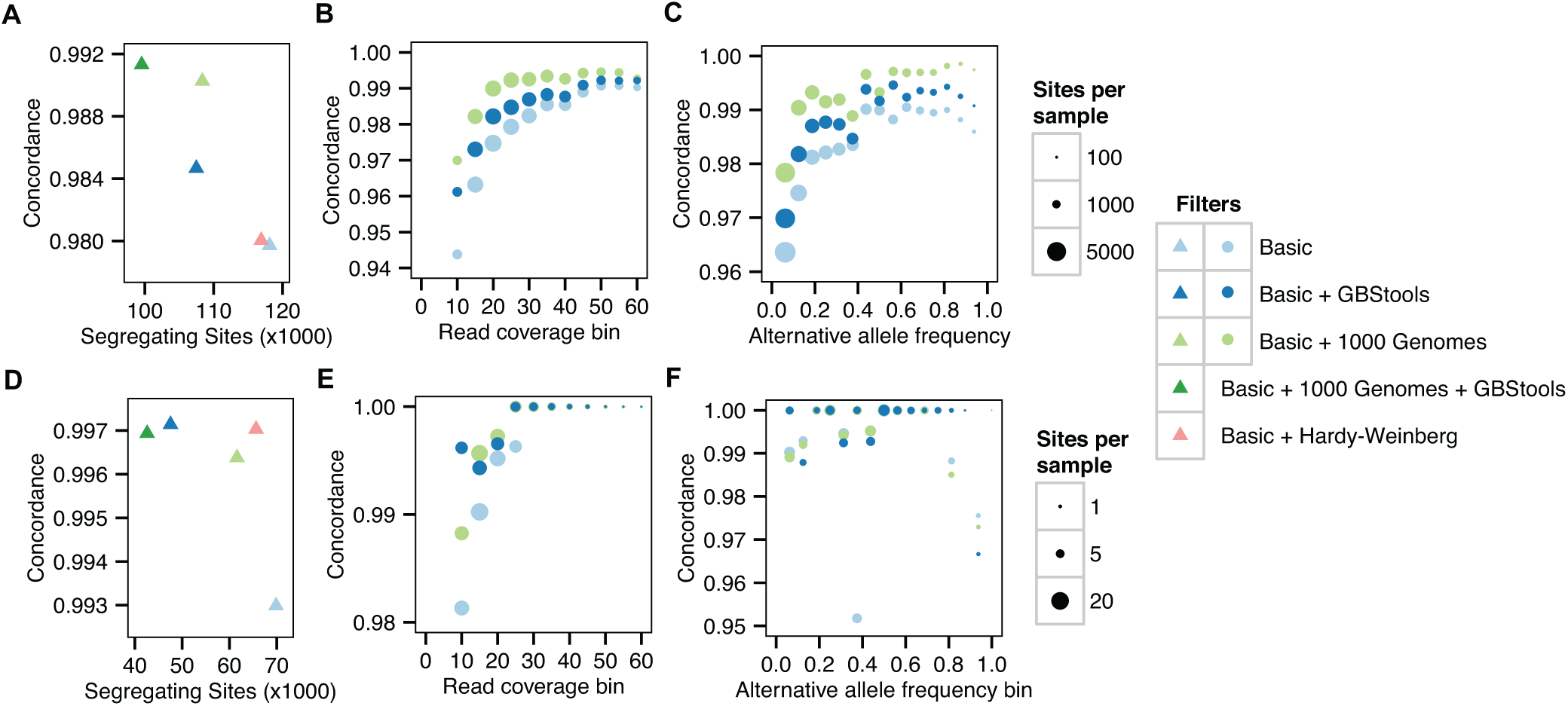
Concordance before and after applying GBS SNP filters. **A.** Proportion of GBS genotype calls concordant with Complete Genomics heterozygote calls for HapMap individuals vs number of segregating sites after applying various filters. Sites passing the basic filters had: mapping quality ≥ 57, SNP quality ≥ 30, coverage ≥ 8X in all samples, and position outside the 1000 Genomes Project callability mask. Sites failing the GBStools filter had: Non-cut allele frequency estimate > 0.05, or likelihood ratio > 2.71 (p < 0.05). Sites failing the 1000 Genomes filter had > 10% of spanning reads mapped to known polymorphic restriction site (allele frequency > 0.01). **B.** Same data as in A, but with genotypes binned by depth of coverage. **C.** Same data as A-B, but with genotypes binned by alternative allele frequencies, which were inferred from whole genome sequencing of the eight HapMap individuals (two sequenced by SOLiD technology, and six sequenced by Complete Genomics). **D-F.** Same analysis as A-C, but for concordance between GBS genotypes calls and exome array calls for the Argentine individuals. Basic filters are same as in A-C, but require ≥ 8X coverage in ≥ 40/63 samples. Allele frequencies were estimated from genotypes of 389 Argentine individuals on the exome array.

The fact that hard filters resulted in a fairly low error rate (2%) suggested that this is a sensible approach for species with genetic diversity similar to humans. But many non-model organisms have higher levels of genetic diversity, which may lead to an error rate that is high enough to necessitate a more sophisticated approach. To explore this possibility, we simulated GBS data under a neutral coalescent model[19] with population mutation rates (θ = 4Nμ) between 1×10^−3^−2×10^−2^. In a preliminary filtering step, we removed SNVs with > 10% missing genotypes, which reduced the genotype error rate to 1.2% for data simulated with θ = 1×10^−3^ (typical of human data), and 4.7% for data simulated with θ = 5×10^−3^, which is typical of high-diversity species such as Drosophila (S4 Fig. A). We simulated 40X GBS coverage for these same genotype data, and found that the GBStools likelihood ratio test reduced the error more than 10-fold, for instance down to 0.3% in the case of the high-diversity (θ = 5×10^−3^) data set (S4 Fig. A). Although normalized site frequency spectra (SFS) were not substantially affected by restriction site polymorphisms (S4 Fig. D-E), errors in the genotypes themselves may cause problems in some applications. In these cases, particularly in studies of high diversity species, GBStools is expected to improve genotyping accuracy more than hard filters.

As a preliminary step in testing the utility of GBStools, it was necessary to confirm the theoretical prediction that samples with one non-cut restriction site allele (restriction site genotype +/−) have on average half the coverage of samples with two intact restriction site alleles (restriction site genotype +/+). To test this, we measured GBS coverage at known polymorphic restriction sites in the HapMap data (Fig. 3). We applied a normalization to account for variation in total read numbers between libraries (methods), and binned the individual sample coverages according to the mean coverage of +/+ samples at each site. Within each bin, we observed two distinct, but overlapping, coverage distributions for samples with restriction site genotypes +/+ and +/−, suggesting that the prediction holds true. The proportion of the +/− distribution that does not overlap the +/+ distribution provides a rough measure of the potential power of a statistical test for restriction site polymorphism based on read coverage, and it is evident from the extensive overlap of the two distributions in the 5-15X and 25-35X bins that higher coverage is necessary to achieve substantial power. If the goal of a particular study were to estimate population-level summary statistics such as Fst, or to map traits in an experimental cross, the added accuracy afforded by such a test might not be worth the extra sequencing effort to achieve > 35X coverage. If the goal, however, were to estimate the site frequency spectrum, then high genotype accuracy would be necessary, and in such cases (e.g. exome sequencing) coverage in the > 35X range is not uncommon. Thus the conditions for high-sensitivity detection of restriction site polymorphisms might already exist in many experimental designs.

**Fig. 3.**
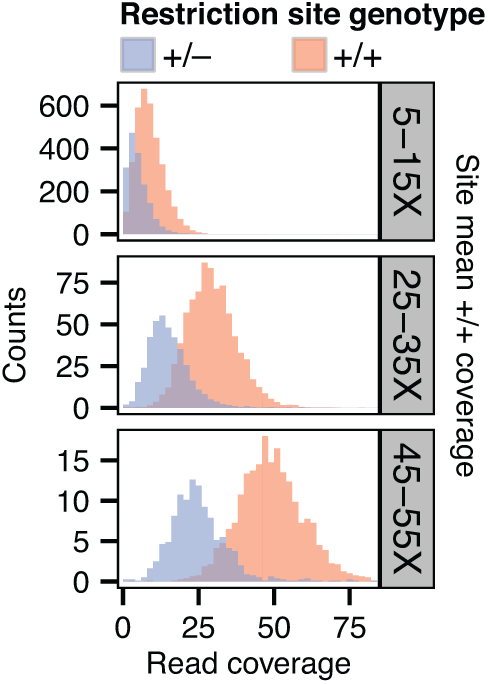
Read coverage distributions at sites with a known restriction site polymorphism. Distributions of normalized depth of GBS coverage for HapMap individuals with one non-cut restriction site allele (genotype +/−) or with two intact restriction site copies (genotype +/+) at sites with a known restriction site polymorphism.

To better define the experimental conditions under which it is possible to use GBStools effectively, we applied GBStools to data simulated with different numbers of samples (from N = 8 to N = 500), and read coverages (10-100X). Since the proportion of homozygotes at a SNV observed by GBS is sometimes inflated by restriction site polymorphism, we also used an exact test to assess the chance of observing the given genotypes (or a worse-fitting set of genotypes) at each site under Hardy-Weinberg equilibrium. We then calculated the sensitivity and specificity of the GBStools likelihood ratio, or the Hardy-Weinberg p-values, as classifiers of incorrect vs correct genotype calls under varying thresholds (Fig. 4A), and measured the area under curve (AUC) of the response operator characteristic (ROC) curves as indicators of the test’s performance. In theory, an uninformative (random) classifier has AUC = 0.5, whereas a perfect classifier has AUC = 1.0. The GBStools test outperformed the Hardy-Weinberg test as measured by area under the curve (AUC), particularly at high-coverage sites (Fig. 4A, S3 Table). We noted that the ROC curves for the GBStools test at low coverage (10X) and the Hardy-Weinberg test have a similar shape, which may be due to the assumption of Hardy-Weinberg genotype proportions in the GBStools model (S1 Appendix). Aside from the already-established benefit of high coverage, we also found that large sample sizes were beneficial to GBStools performance. For example, power to detect non-cut restriction site alleles of frequencies between 0.01-0.02 was 25% for 30 samples at 40X coverage, but was 94% for 500 samples at the same coverage (Fig. 4B). For 40X sites in the 100- and 500-sample data sets, AUC was at least 0.96, suggesting that this is the ideal coverage and sample size range for using GBStools.

**Fig. 4.**
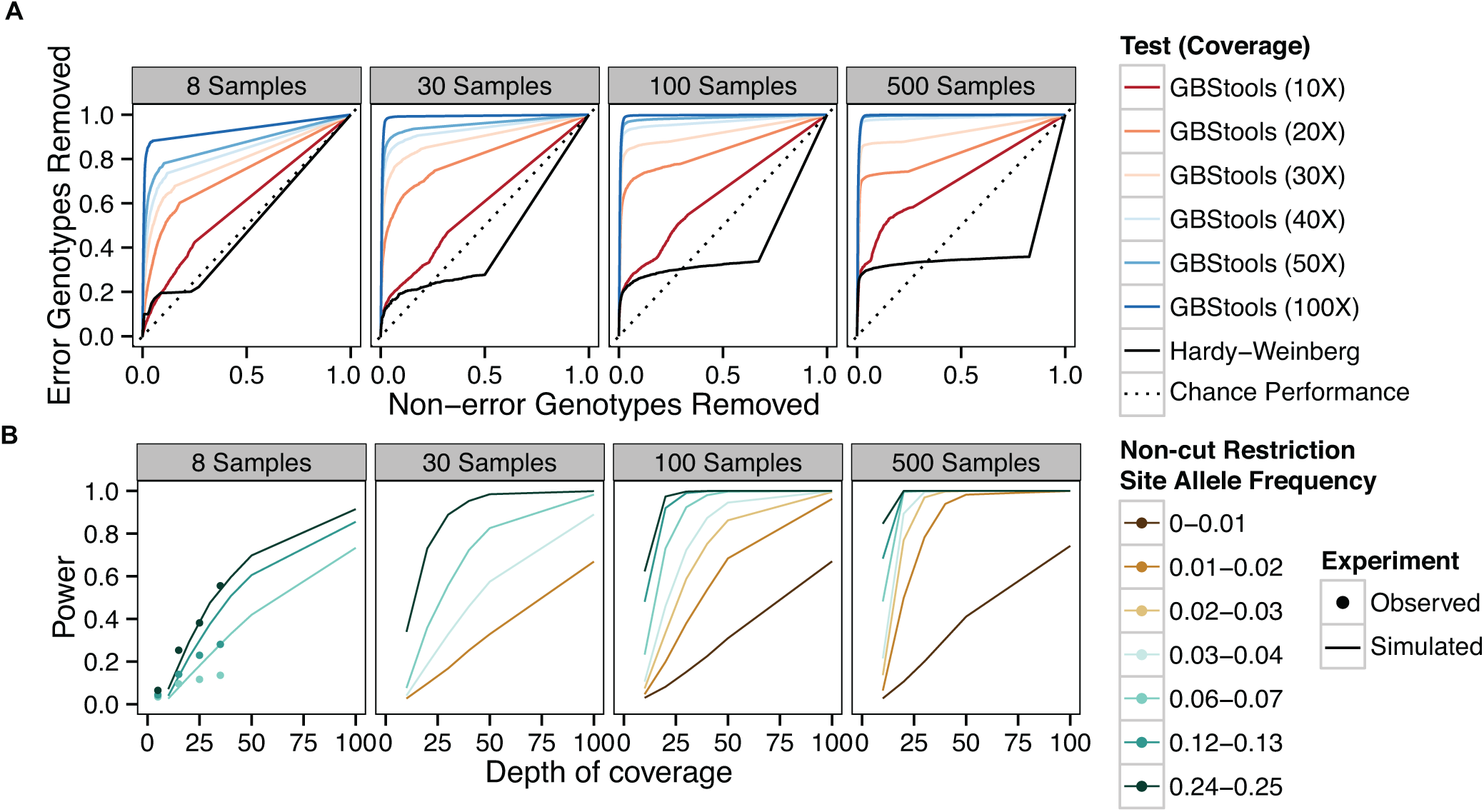
Sensitivity and specificity of GBStools likelihood ratio test. **A.** Response operator characteristic (ROC) curves for classification of incorrect vs correct heterozygote genotype calls by GBStools likelihood ratio test or Hardy-Weinberg equilibrium exact test p-values. The data were filtered by call rate (sites with > 10% missing genotypes were excluded) before applying either test, and the axes refer to the proportion of genotypes that passed this filter. The diagonal represents performance of an uninformative (random) classifier. **B.** Power of GBStools likelihood ratio test for detecting restriction site polymorphism with simulated and empirical data. We used a critical value of 2.71 for calculating power, based on the expected null distribution, a one-half chi-squared distribution with one degree of freedom (p < 0.05). Empirical power was calculated for 331,861 autosomal SNPs in the HapMap GBS data set that passed insert size filters and where the EM parameter estimates converged (see Methods).

At lower coverage (10-20X) and with smaller sample sets (N = 8) GBStools did not perform as well in simulations (Fig. 4), and this may explain the modest increase in concordance from 98.0% to 98.5% when the GBStools filter was applied to the HapMap data set (N = 8), which led to the removal of 9% of segregating sites (Fig. 2A). For comparison, a filter for known restriction site polymorphisms in the 1000 Genomes Project[18] data set improved concordance to 99.0% (Fig. 2A, S2 Table C), suggesting that the power of GBStools was no higher than 50%. Indeed, power to detect common restriction site polymorphisms in the HapMap GBS data (non-cut allele frequency 0.25) was 56% for sites covered to 30-40X, but for singleton sites covered to 30-40X it was only 13%, which was lower than predicted by simulation (Fig. 4B). In addition, AUC values for the HapMap ROC curves were lower than the values obtained in simulations with matching coverage levels (Fig. 5A, S3 Table). This is possibly due to the model’s assumption of a constant value for the index of dispersion in depth of coverage between samples, whereas the empirical data exhibit variation in dispersion from site to site (S5 Fig.). It should be possible to relax this assumption by estimating dispersion on a site-by-site basis, or by calculating a joint estimate from genome-wide data, but these methods are currently not implemented. Joint modeling of genotypes at multiple closely-linked SNPs should also offer an increase in power over the single-marker model currently implemented. This would be particularly useful in the case of long reads, where each “stack” of reads mapped to a particular restriction site would contain more SNPs on average than a stack of shorter reads. For the present time, however, our simulations suggest the easiest way to improve the low empirical power observed here is to increase the number of samples.

**Fig. 5.**
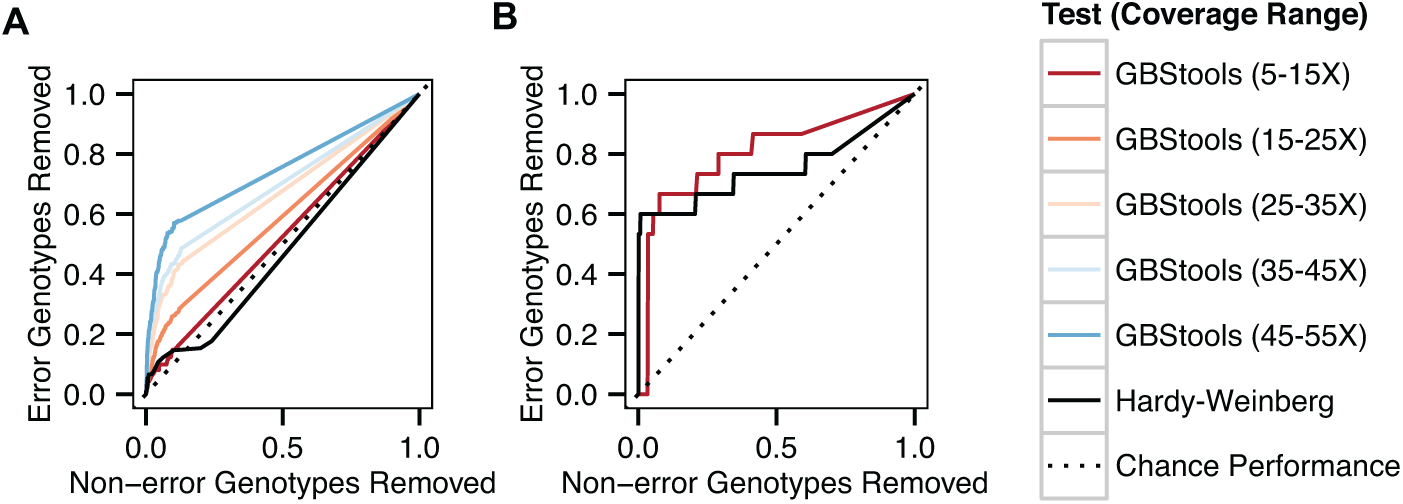
Empirical sensitivity and specificity of GBStools test. **A.** Response operator characteristic (ROC) curves, as in Fig. 4A, for GBStools and Hardy-Weinberg tests with HapMap data. The data were filtered by coverage, call rate, and mapping quality (methods), before applying either test. **B.** Same as (A), but for Argentine data set.

We investigated whether it is possible to accurately estimate which particular genotypes are likely to be affected by allelic dropout. As mentioned in the introduction, the true numbers of non-cut alleles per sample are latent variables in the GBStools likelihood model, and the expected values of these variables are output by GBStools in VCF format. We compared these expected non-cut allele counts to the true counts inferred from whole-genome sequencing data to gain an idea of their predictive value (Fig. 6). Although samples with a true allele count of one (i.e. restriction site genotype +/−) had higher average expected non-cut allele counts than samples with true allele count of zero (genotype +/+), it is clear that this is not a very sensitive predictor. For instance, +/− samples at sites with non-cut allele frequency 0.25 and 30-40X coverage had a median expected non-cut allele count of 0.01 (Fig. 6), far from the true value of 1.0. Yet power to detect restriction site variants in these same data was 56% (Fig. 4B). This indicates that the true utility of GBStools is in determining whether or not any samples at a site carry non-cut alleles rather than determining which particular samples carry them, although in some cases (Fig. 1C) there is diagnostic value in the latter approach.

**Fig. 6.**
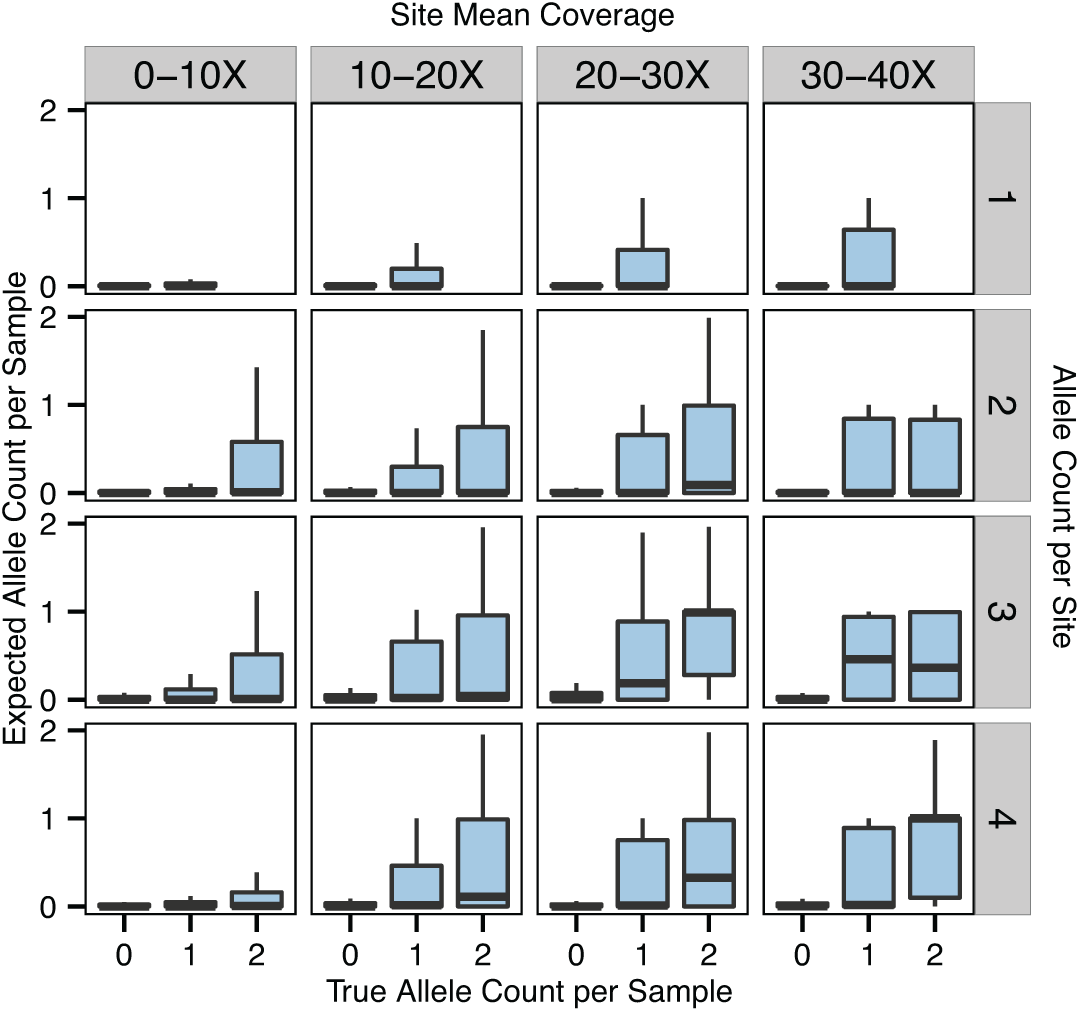
Expected non-cut restriction site allele counts. The true numbers of non-cut restriction site alleles carried by each sample are latent variables in the GBStools likelihood model. Boxplots representing the distributions of the expected values of these variables are shown here, and are grouped by the true non-cut allele counts inferred from Complete Genomics whole-genome sequencing data for HapMap samples. Allele counts of 0, 1 and 2 correspond to restriction site genotypes +/+, +/− and −/− respectively, where (−) is the non-cut allele. Plots are also grouped by site allele count (with 8 samples total) and site mean coverage. Only sites with allele frequencies between 0.0625–0.25 are shown.

The site frequency spectrum derived from our filtered GBS data was similar to the spectrum from whole-genome sequencing data, with 2.3% fewer singletons (Fig. 7A). This suggested that GBS data can be useful in population genetic studies, for example demographic inference based on the site frequency spectrum.

**Fig. 7:**
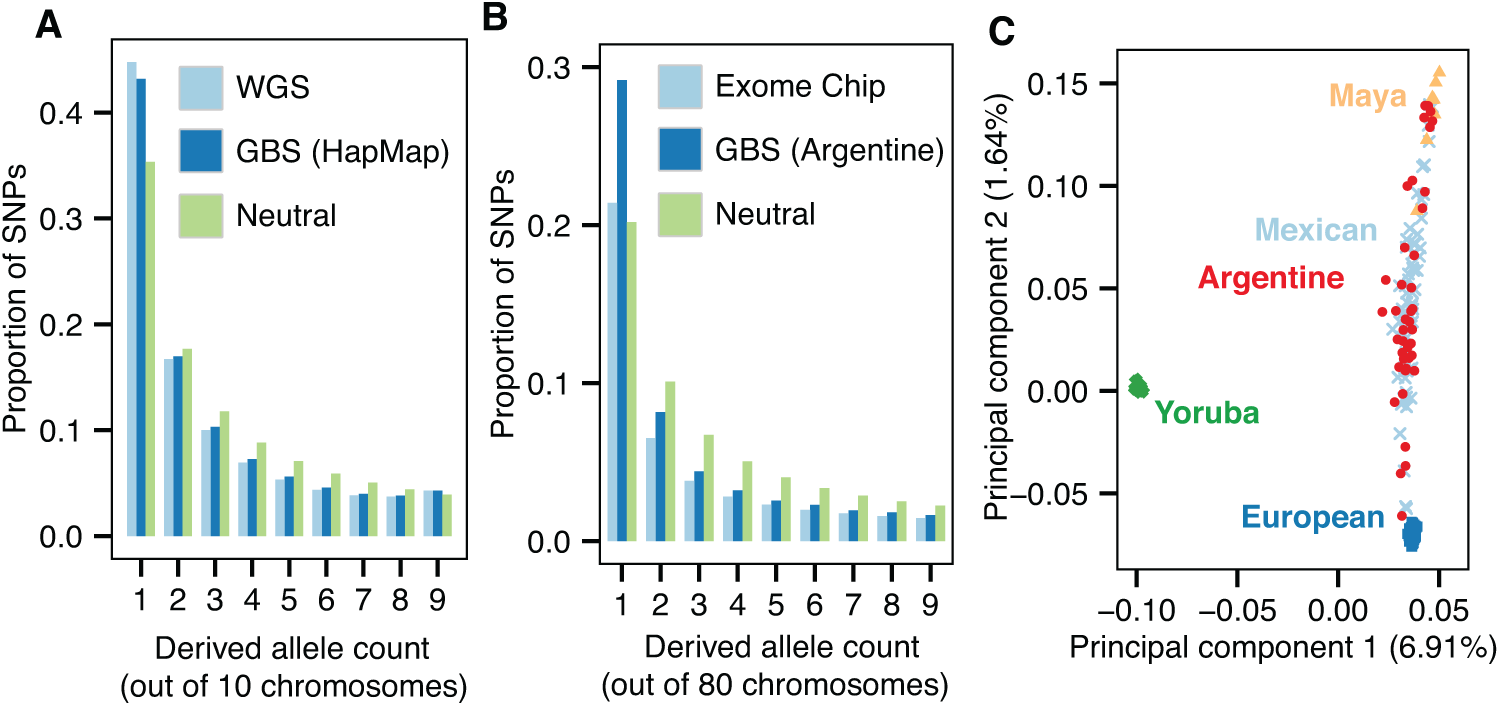
Detecting population structure and growth with GBS data. **A.** Normalized site frequency spectra (SFS) of the derived allele for six HapMap samples represented as the expected SFS of a subsample of size five to account for missing data. SNPs from the 29.2 Mb region that passed all filters were used for both GBS and whole-genome sequencing (WGS) spectra. The expected SFS under a neutral coalescent model is shown for comparison. **B.** Bins 1–9 of the normalized SFS for SNPs in the Argentine GBS data set, represented as the expected SFS of a subsample of size 40. Exome chip data was from 386 Argentine individuals, including some of those sequenced by GBS. **C.** Principal components 1 and 2 of the admixed Argentine individuals, Europeans (CEU), Yoruba (YRI), Mexican (MXL), and Maya (HGDP), using 30,691 SNPs. Of the Argentine samples, 40/63 passed the 5% data missingness filter and were used in the PCA.

To explore this further, we sequenced 89 admixed Argentine individuals to test for signatures of mixed ancestry and demographic changes (S4 Table). The Argentine samples had 7.5X mean coverage in a 177 Mb target region (S3 Fig. B, S4 Table). Argentine samples with < 30% of reads mapped to restriction sites (26/89 samples) were excluded from further analyses, as it is likely that these samples were not digested to completion. A total of 1,013,785 segregating sites were called in the remaining samples and concordance with exome array data was 99.7% after filtering with GBStools, which led to removal of 25% of sites (Fig. 2D, S3 Table H-K). A filter for Hardy-Weinberg equilibrium showed similar sensitivity and specificity (Fig. 5B), although fewer segregating sites were removed (Fig. 2D), indicating the GBStools critical value we used was more conservative. Both tests performed better than expected in simulations with a similar number of samples (N = 100). This is probably due to the small number of errors that remained after applying basic filters (15 in total, see S3 Table), and the fact that over half of these errors originated from a single SNP (rs6861689) that is near a common restriction site polymorphism (BsaXI site overlapping rs6861731).

We calculated the expected SFS from the Argentine GBS data and compared it to the SFS under a neutral coalescent model, and to the SFS from 386 Argentine individuals genotyped on an exome SNP array (Fig. 7B). The excess of singletons in the GBS spectrum is consistent with recent population growth,[20] but was not observed in the array data, most likely due to ascertainment bias.

Another potential area where GBS can be useful is in ancestry estimation. We joined the Argentine GBS data set with SNP data from Yoruban, European, and Mexican individuals from the 1000 Genomes Project[18] phase 1 data set, and from Mayan individuals from the Human Genome Diversity Project, and performed principal components analysis (Fig. 7C, methods). As expected, individuals from the admixed Argentine populations fell between the European and Native American populations in PC space.

In summary, we have used high-quality human SNP chip and whole-genome sequencing resources to test several different methods for reducing genotype errors in GBS data, including commonly-used hard filters, and a new GBS-specific statistical method implemented in our open-source program GBStools. These methodological improvements enable GBS to nearly match whole-genome sequencing in accuracy, as we have demonstrated, but at a fraction of the cost. Furthermore, our simulations suggested that GBStools has substantially better performance than hard filters in high diversity species with extensive restriction site polymorphism. Since GBStools is designed to accept data in the standard VCF format (and can optionally use read data in the standard SAM/BAM format), it can supplement many pre-existing GBS variant calling pipelines, for example the one implemented in the program Stacks[21]. We anticipate that this approach may enable many GBS-based analyses beyond high-throughput trait mapping, in particular population genetics studies such as detecting signatures of hitchhiking and selection, and estimating demographic history.

## Methods

### Simulation of GBS data

We used Hudson’s ms[19] to generate 1×10^7^ random samples of 200 haplotypes at a 500 bp-long locus with a population mutation rate of 1×10^−3^ (θ = 4N_e_μ) without recombination. The position of each segregating site within the locus was drawn from a uniform distribution. The first and last 6 bp of the locus represented two 6 bp-long restriction enzyme recognition sites. If any segregating site fell within these two sites, a restriction site polymorphism resulted, and either the derived or ancestral allele was randomly chosen to represent the non-cut restriction site allele. Segregating sites in the interior of the fragment, but farther than 6 bp from the ends, were chosen to represent restriction site polymorphisms with probability 0.0074 (the frequency of bases that are part of BpuEI, BsaXI, and CspCI recognition sites in the human genome). Segregating sites within 101 bp of the fragment ends represented sites sequenced by GBS with paired-end 101 bp reads. We randomly paired the 2N haplotypes to create a set of N diplotypes. Heterozygous genotypes within the ‘read’ portion of diplotypes that were heterozygous for one of the restriction sites were counted as genotyping errors. Simulations with population mutation rates of 5×10^−3^, 1×10^−2^, and 2×10^−2^ were also carried out. As most loci simulated in this manner do not carry restriction site polymorphisms it is an inefficient way to simulate large numbers of them. Thus to simulate GBS data for estimating the power of the GBStools likelihood ratio test we randomly chose one segregating site per locus to represent a restriction site polymorphism, irrespective of its location, and randomly chose either the derived or ancestral allele to be the non-cut allele. Depth of coverage was drawn from a negative binomial distribution with mean μ and scale parameter μ / 1.5 (dispersion index = 2.5). Read likelihoods were then calculated[17], assuming a constant sequencing error rate of 1×10^−3^.

### Genomic DNA Samples

Genomic DNA from eight HapMap individuals, including six samples sequenced by Complete Genomics[15] and two samples sequenced with SOLiD technology[16], was obtained from Coriell Cell Repositories. The Argentine samples were collected from 15 geographical regions in Argentina in multiple sampling efforts between 2007-2012. Under local IRB approval, blood samples were collected from participants who gave informed consent. Both HapMap and Argentine samples were de-identified and analyzed anonymously. Blood was stored in lysis buffer[22] in the field before genomic DNA was extracted in the lab by a standard salting-out procedure[23].

### Exome array genotype calls

Data for Illumina Human Exome Beadchip v1.0 (HumanExome-12v1_A) were generated for the Argentine samples at the Hussman Institute for Human Genomics, University of Miami. Genotypes were called with Illumina’s Genome Studio V2011.1 with a no-call threshold of 0.15. A minimum call rate of 99.3% was required for each sample and 386 of the 391 Argentinean samples passed this filter. Per-SNP quality filters included: mapping to a unique genomic location, and minimum per-SNP call rate of 99% (245,937 SNPs met these criteria). Of these sites, 8 were excluded from the concordance analysis for the reason that more than one sample had an exome array call of homozygous reference and a GBS call of homozygous non-reference (or vice versa).

### Whole genome sequencing variant calls

Variation data files (masterVar) for samples NA18505, NA18508, NA19648, NA19704, NA21732, and NA21733 were downloaded from the Complete Genomics ftp site. We generated a vcf file with the mkvcf utility (v1.6.0 build 43). Before calculating concordance with GBS calls, we removed low confidence and hemizygous genotype calls, and excluded 10 sites that exhibited discordance with the GBS calls across the majority of samples. We used the unfiltered variant calls for site frequency spectrum estimation, but split multi-nucleotide polymorphisms into their component SNPs with a custom python script. We used another custom python script to predict BpuEI, BsaXI, and CspCI restriction site variants caused by bi-allelic SNPs and indels in the unfiltered calls. The sequencing of samples NA19740 and NA19836 was described previously[16]. We predicted restriction site polymorphisms caused by SNPs in these samples in the same manner.

### Library Preparation

Genomic DNA (50 ng) was digested with BpuEI (2.5 U), BsaXI (2 U), and CspCI (2.5 U) (NEB) at 37° for 90-120 min in buffer containing 20 μM S-adenosylmethionine. The digestion product was purified on a DNA Clean and Concentrate column (Zymo Research). DNA end repair, 3’ monoadenylation, and ligation of sequencing adapters were performed as described in the Illumina TruSeq DNA Sample Preparation Guide. We designed a custom set of sequencing adapters, derived from the TruSeq adapters, with 65 six-bp barcodes (S5 Table). We used a standard protocol to anneal the common adapter to each of the 65 barcode adapters[8]. The ligation product was amplified by 10 cycles of PCR. For the HapMap samples, inserts between 350-650 bp were size selected on a Caliper Labchip, with one sample per gel lane. For the Argentine samples, inserts between 350-650 bp were size-selected in batches of 9-11 samples per gel lane. Bioanalyzer quantification was used to pool in equimolar amounts before and after size selection. For the 89 Argentine samples, two pools were prepared and sequenced separately, the first with 24 samples and the second with 65. Because of the high variance in read numbers per sample we observed in the Argentine libraries, we later re-analyzed the bioanalyzer data from the first set of 24 samples (S1 Fig.).

### Sequencing and read mapping

Libraries were sequenced on the Illumina HiSeq 2000 in 2 × 101 bp mode following the standard TruSeq SBS protocol. The eight HapMap samples were sequenced on a single lane, with a mean of 18.3 M paired end reads per sample. In the Argentine study, the two pooled libraries were sequenced on four and five separate lanes respectively, with a mean of 17.5 M reads per sample. Reads were mapped to the human reference genome (build 37) with BWA[24] with the -q 20 parameter to include soft clipping of low quality bases. Local realignment of reads around known indels and base quality recalibration were performed with GATK[17]. We defined the target region for the HapMap samples by taking the union of predicted restriction site fragments between 400-700 bp that had ≥ 3X mean coverage, and where ≥ 10% of reads had a mate pair mapped to a restriction site (S2 Fig. A). The target region for the Argentine samples was defined in the same way, but with predicted fragments between 200-600 bp. Argentine samples with < 30% of reads mapped to restriction sites (26/89 samples) were excluded from further analyses.

### Calculation of coverage distributions at polymorphic restriction sites

For each of the HapMap samples in our GBS data set we inferred the number of cut and non-cut alleles at each restriction site in the genome from the Complete Genomics and SOLiD data. We then calculated depth of coverage and median insert size at each site. For this analysis we kept only sites where the median insert sizes were between 350-625 bp for each sample, and where ≤ 4 samples had zero depth of coverage. We normalized the depth of coverage for each sample by multiplying by the following normalization factor:

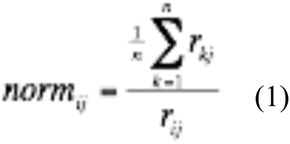

Here *n* is the total number of samples, and *r_ij_* is the total number of library inserts of size *j* for individual *i*. In calculating *norm_ij_* for a particular site we took *j* to be the median insert size of reads from individual *i* at that site. We then binned each site according to the mean coverage of samples that had two restriction site copies. Then, aggregating the coverage data across samples, we plotted the coverage distributions for each bin.

### Variant calls and hard filters

We called SNPs in the target regions described above with the GATK Unified Genotyper, emitting both variant and invariant sites. We also used the GATK Haplotype Caller to call SNPs in the HapMap data set. We found that specificity was higher for Haplotype Caller, with fewer true homozygous reference genotype called heterozygous (S2 Table F), but also found that sensitivity was lower, with fewer true SNPs called. It is possible that this was because we used Haplotype Caller parameters that are optimal for whole-genome sequencing but not for GBS. We did not explore this point further, however, and instead used the SNP calls from Unified Genotyper for the remainder of the analyses. We performed variant quality score recalibration on segregating sites with GATK with the following training data sets (downloaded from the Broad Insitute ftp server): hapmap_3.3.b37.sites.vcf 1000G_omni2.5.b37.sites.vcf. For the HapMap samples we also trained with known variants from previous whole-genome sequencing studies[15,16]. We trained VQSR with the annotations HaplotypeScore, QD, ReadPosRankSum and HRun, and kept sites in the 99% sensitivity tranche. Invariant sites were not subjected to the VQSR filter. We applied the following hard filters (labeled as ‘basic filters’ in figures and tables): mapping quality ≥ 57, SNP quality ≥ 30, coverage ≥ 8X in all samples (HapMap samples) or coverage ≥ 8X in ≥ 40/63 of samples (Argentine samples). We also filtered out sites that fell within the 1000 Genomes Project callability masks for depth of coverage and mapping quality. In addition, we applied a filter for sites where the observed genotypes differ significantly from those predicted under Hardy-Weinberg equilibrium (p < 0.05), with the software package vcftools[25].

### 1000 Genomes Project polymorphic restriction site filter

We used a custom python script to predict BpuEI, BsaXI, and CspCI restriction site variants caused by SNPs and indels in the 1000 Genomes Project data set. For each sample we created a set of genomic intervals where more than five read pairs spanned a restriction site that was polymorphic with a minor allele frequency of > 0.01. We then filtered out all sites that fell within the interval set of more than one sample.

### GBStools polymorphic restriction site filter

The calculation of frequency estimates for non-cut restriction site alleles, and the calculation of the likelihood ratio test statistic for restriction site polymorphism are described in S1 Appendix. We implemented these algorithms in the python package GBStools (http://med.stanford.edu/bustamantelab/software.html). Frequency estimates for a non-cut restriction site allele are expected to be zero under the null hypothesis (no polymorphism). Since this is on the boundary of the parameter space (0, 1], the parameter estimate is expected to have a half-normal distribution. Therefore, the test statistic is expected to have an approximately one-half chi-squared distribution with one degree of freedom[26], which has a critical value of 2.71 (p = 0.05). We applied the likelihood ratio test to simulated GBS data and found that at high coverage the test statistic was equal to zero more often than expected (S6 Fig.). In the 20-50X coverage range, however, it agreed well with the expected distribution. The departure from the expected null distribution at high coverage was related to the fact that more than half of the allele frequency estimates were zero (S7 Fig.) and suggested that in general 2.71 is a lenient critical value (p < 0.05) for detecting restriction site polymorphisms. We performed the likelihood ratio test for SNPs where the median insert size was between 450-625 bp (HapMap individuals) or 300-500 bp (Argentine individuals) and where the median absolute deviation in insert size was less than 60 bp (S8 Fig.). For the ‘GBStools filter’ listed in the figures and tables, we kept only SNPs that had a likelihood ratio < 2.71 and an estimated frequency of the non-cut restriction site allele < 0.05. In addition, we excluded the region spanned by the two restriction sites nearest to any site that did not meet these criteria.

### GBStools power calculation

We applied the likelihood ratio test described above to GBS data from the HapMap samples. We restricted the power analysis to autosomal sites that were segregating in the Complete Genomics data set, where the median GBS insert size was between 450-625 bp, and the median absolute deviation for insert sizes was ≤ 60 bp (331,861 sites). We binned the sites according to mean depth of coverage, and for each bin we calculated the power to detect known polymorphic restriction sites at a conservative critical value of 2.71 (empirical p = 0.05 critical values were slightly lower).

### Site frequency spectra

We calculated the expected site frequency spectrum from GBS data and Complete Genomics data for HapMap samples NA18505, NA18508, NA19648, NA19704, NA21732, and NA21733 as a subsample of size five in order to allow for missing data[27,28]. We used 1000 Genomes inferred ancestral alleles, and discarded sites where the ancestral allele was not consistent with the observed alleles. We kept sites that passed variant quality score recalibration and passed the hard filters (‘basic filters’), the 1000 Genomes Project restriction site polymorphism filter, and the GBStools filter (29.2 Mb of total unmasked sites). The whole-genome sequencing (Complete Genomics) site frequency spectrum was calculated based on segregating sites in this same region. We calculated the expected site frequency spectrum for the Argentine samples as a subsample of size 40 after applying the filters shown in S2 Fig. (12.7 Mb of total unmasked sites). We also calculated the expected site frequency spectrum for 386 Argentine individuals genotyped on the Illumina exome chip, as described above. We used exome chip genotypes located in both filtered and unfiltered regions.

### Principal components analysis

We merged the Argentine GBS data with 1000 Genomes Project SNP data (CEU, YRI, and MXL populations), and with HGDP SNP data from sequenced Mayan individuals[29]. Of the segregating sites in the merged data set, 715,082 were present in each of the original data sets. We kept Argentine individuals that had > 25% of these sites sequenced to ≥ 7X (42/63 samples were kept). We then filtered out sites where < 90% of all samples had called genotypes. We then applied the hard filters listed previously, and pruned SNPs for linkage disequilibrium (r^2^ < 0.8 in 50 bp windows with 5 bp step size) with PLINK[30], resulting in a final set of 45,630 SNPs. We performed principal components analysis on this set of SNPs with smartpca[31].

## Acknowledgements

The authors thank Suyash Shringarpure (Bustamante lab) for helpful suggestions, Brenna Henn and Jeffery Kidd for early access to the Mayan sequencing data, and Josefina MB Motti, Laura S Jurado-Medina, M Rita Santos, Virginia Ramallo, Marisol Schawb, Julieta Beltramo, and Mariela Cuello for help in collecting DNA samples, and the following groups from the University of Miami: Hussman Institute for Human Genomics, Center for Genome Technology, Genotyping Core, Biorepository Core.

